# FORMATION OF MALIGNANT, METASTATIC SMALL CELL LUNG CANCERS THROUGH OVERPRODUCTION OF cMYC PROTEIN IN TP53 AND RB1 DEPLETED PULMONARY NEUROENDOCRINE CELLS DERIVED FROM HUMAN EMBRYONIC STEM CELLS

**DOI:** 10.1101/2023.10.06.561244

**Authors:** Huanhuan Joyce Chen, Eric E. Gardner, Yajas Shah, Kui Zhang, Abhimanyu Thakur, Chen Zhang, Olivier Elemento, Harold Varmus

## Abstract

We recently described our initial efforts to develop a model for small cell lung cancer (SCLC) derived from human embryonic stem cells (hESCs) that were differentiated to form pulmonary neuroendocrine cells (PNECs), a putative cell of origin for neuroendocrine-positive SCLC. Although reduced expression of the tumor suppressor genes *TP53* and *RB1* allowed the induced PNECs to form subcutaneous growths in immune-deficient mice, the tumors did not display the aggressive characteristics of SCLC seen in human patients. Here we report that the additional, doxycycline-regulated expression of a transgene encoding wild-type or mutant cMYC protein promotes rapid growth, invasion, and metastasis of these hESC-derived cells after injection into the renal capsule. Similar to others, we find that the addition of cMYC encourages the formation of the SCLC-N subtype, marked by high levels of *NEUROD1* RNA. Using paired primary and metastatic samples for RNA sequencing, we observe that the subtype of SCLC does not change upon metastatic spread and that production of NEUROD1 is maintained. We also describe histological features of these malignant, SCLC-like tumors derived from hESCs and discuss potential uses of this model in efforts to control and better understand this recalcitrant neoplasm.

## Introduction

From the earliest days of cancer research, attempts to understand the mechanisms of carcinogenesis have depended on experimental models to circumvent the ethical obstacles to performing experiments in human subjects. Such model systems have ranged from normal and cancerous human cells grown in animals or in cell culture to the generation or manipulation of cancers arising from normal cells in whole animals. The success of such approaches is often measured by the similarities of model systems to the observed traits of cancers that have arisen naturally in human subjects. But the utility of certain model systems can also be appraised by evaluation of the information that emerges from them, even when the models lack a clear relationship to any specific type of human cancer. The transformation of chicken embryo fibroblasts by Rous sarcoma virus or the induction of leukemias with various murine leukemia viruses exemplify models that lack verisimilitude but nevertheless have revealed major features of carcinogenesis (1).

A common complaint about many models of cancer is that the transformation of a normal cell to a malignant cell occurs in non-human cells. Efforts to transform human cells are often limited by the difficulties of growing human cells in culture and uncertainty about the developmental stage of a given cell lineage at which transformation generally occurs. Techniques for generating, propagating, and differentiating human stem cells - either tissue-specific stem cells or pluripotent stem cells - have raised the possibility of studying carcinogenesis in a wide variety of human cell lineages, as initially demonstrated for gliomas derived from induced human pluripotent stem cells (iPSCs) (2).

We have recently attempted to study the mechanisms by which small cell lung cancer (SCLC) arises in human lung cells by efficiently differentiating human embryonic stem cells (hESCs) (3, 4) into mature lung epithelial cells, including the likely precursor of SCLC (pulmonary neuroendocrine cells [PNECs]) and then simulating loss of function of the tumor suppressor genes *TP53* and *RB1*, tumor suppressors commonly lost in human SCLC (5). PNEC-like cells that have lost the function of these two tumor suppressor genes, by expression of short hairpin RNAs (shRNAs) specific for *RB1* and *TP53*, can form tumors subcutaneously in immunodeficient mice; however, these tumors grow slowly and do not invade or metastasize, standing in contrast to what is known about the clinical presentation and behavior of SCLC (6). Thus, the original hESC-derived SCLC models we generated appeared to be benign growths that do not resemble SCLCs functionally, limiting their utility as models for aggressive human SCLC.

Our earlier findings are consistent with the idea that full transformation of PNECs to form malignant SCLC requires additional changes beyond restricted expression of the tumor suppressor genes *RB1* and *TP53*, such as heightened expression of a proto-oncogene in the *MYC* family (7, 8). In an attempt to show that we can generate more aggressive, SCLC-like tumors from PNECs derived from hESCs, we have added efficiently transcribed *cMYC* oncogenes to cells in which expression of *TP53* and *RB1* are inhibited by shRNAs. When these cells were then injected into renal capsules of immunodeficient mice, some produced rapidly expanding tumors that invaded locally and sometimes formed metastatic growths, mostly in the liver.

In this report, we describe morphological and transcriptional features of these tumors and discuss how the addition of putative drivers of SCLCs can be used to study the mechanism of carcinogenesis and to test candidate therapies for this “recalcitrant” cancer.

## Materials and Methods

### Generation of lung cells, including PNECs, from hESCs

Protocols for maintenance of hESCs and generation of pulmonary neuroendocrine cells (PNECs), and other lung cells were modified from previous studies (3-5). Two hESC lines, RUES2 (Rockefeller University Embryonic Stem Cell Line 2, NIH approval number NIH hESC-09-0013, Registration number 0013; passage 7-10) and ES02 (HES2, NIH registry, WiCell Research Insititute, Inc. passage 3-7) were cultured and maintained in an undifferentiated state on irradiated mouse embryonic fibroblasts (Global Stem, cat. no. GSC-6001G). All cells were purchased from ATCC or WiCell in the past 3 years and tests for mycoplasma were negative. The hESC lines were regularly checked for chromosome abnormalities and maintained with normal chromosome numbers. All embryonic stem cell studies were approved by the Institutional Review Board (IRB) at the University of Chicago, or by the Tri-Institutional ESCRO committee under protocols 2016-01-002 and 2017-10-004 (Weill Cornell Medicine, Memorial Sloan Kettering Cancer Center, and Rockefeller University).

hESC differentiation into endoderm was performed in serum-free differentiation (SFD) media of DMEM/F12 (3:1) (Life Technologies) supplemented with N2 (Life Technologies), B27 (Life Technologies), ascorbic acid (50 μg/ml, Sigma), Glutamax (2 mM, Life Technologies), monothioglycerol (0.4 μM, Sigma), 0.05% bovine serum albumin (BSA) (Life Technologies) at 37C in a 5% CO_2_ / 5% O_2_ / 90% N_2_ environment. hESCs were treated with Accutase (Stemcell technology) and plated onto low attachment 6-well plates (Corning Incorporated, Tewksbury MA), resuspended in endoderm induction media containing Y-27632 (10 μM), human BMP4 (0.5 ng/ml), human bFGF, (2.5 ng/ml), and human Activin A (100 ng/ml) for 72 – 84 hours (hrs) dependent on the formation rates of endoderm cells. On day 3 or 3.5, the embryoid bodies were dissociated into single cells using 0.05% Trypsin/0.02% EDTA and plated onto fibronectin-coated (Sigma), 24-well tissue culture plates (∼100,000– 150,000 cells/well). For induction of anterior foregut endoderm, the endoderm cells were cultured in SFD medium supplemented with 1.5 μM Dorsomorphin dihydrochloride and 10 μM SB431542 for 36-48 hrs, and then switched to 36-48 hrs of 10 μM SB431542 and 1 μM IWP2 treatment.

For induction of early-stage lung progenitor cells (days 6–15), the resulting anterior foregut endoderm was treated with CHIR99021 (3 μM), human FGF10 (10 ng/ml), human FGF7 (10 ng/ml), human BMP4 (10 ng/ml) and all-*trans* retinoic acid (ATRA; 50nM) in SFD medium for 8–10d. Day 10–15 cultures were maintained in a 5% CO2/air environment. On days 15 and 16, the lung field progenitor cells were re-plated after brief one-minute trypsinization onto fibronectin-coated plates, in the presence of SFD containing either a combination of five factors: CHIR99021 (3 μM), human FGF10 (10 ng/ml), human FGF7 (10 ng/ml), human BMP4 (10 ng/ml), and ATRA (50 nM), or three factors: CHIR99021 (3 μM), human FGF10 (10 ng/ml), and human FGF7 (10 ng/ml) for days 14-16. Day 16–25 cultures of late-stage lung progenitor cells were maintained in SFD media containing CHIR99021 (3 μM), human FGF10 (10 ng/ml), and human FGF7 (10 ng/ml) in a 5% CO2 / air environment.

For differentiation of mature lung cells (days 25–55), cultures were re-plated after brief trypsinization onto 3.3% matrigel coated 24-well plates in SFD media containing maturation components CHIR99021 (3 μM), human FGF10 (10 ng/ml), human FGF7 (10 ng/ml), and DCI (50 nM Dexamethasone, 0.1 mM 8-bromo-cAMP and 0.1 mM IBMX (3,7-dihydro-1-methyl-3-(2-methylpropyl)-1*H*-purine-2,6-dione). DAPT (10 µM) was added to the maturation media for induction of PNECs. All growth factors and most small molecules used for hESC differentiation were purchased from R&D Systems. ATRA, 8-bromo-cAMP, IBMX and DAPT were purchased from Sigma-Aldrich.

### Lentivirus transduction of hESCs

HSC lines were constructed to contain DOX-inducible cassettes encoding shRNAs specific for the *TP53* and *RB1* tumor suppressor genes (RP cells) and RP cells containing DOX-inducible cassettes encoding wildtype human cMYC or a stable, mutant version of cMYC (T58A), to generate RPM and RPM (T58A) cells. The lentiviral vector expressing tetracycline (TET)-inducible shRNA against human P53 was purchased from Gentarget. Inc. (cat# LVP-343-RB-PBS). The lentiviral vectors expressing TET-inducible shRNAs against human *RB1* construct (“pSLIK sh human Rb 1534 hyg” was a gift from Julien Sage; plasmid # 31500) (9), TET-inducible wild type cMYC (FUW-tetO-hMYC was a gift from Rudolf Jaenisch; plasmid # 20723) or mutant cMYC (T58A) tagged at the N-terminus with three copies of a hemagglutinin tag (3X-HA) (pLV-tetO-myc T58A was a gift from Konrad Hochedlinger; plasmid # 19763) were obtained from Addgene and sequence verified prior to use. The shRNA target sequences are as follows: RB1 #1: 5’-GGACATGTGAACTTATATA-3’, RB1 #2: 5’-GAACGATTATCCATTCAAA-3’, p53: 5’-CACCATCCACTACAACTACAT-3’.

To generate lentiviral particles, the above plasmids were transfected into HEK293T cells with the PC-Pack2 lentiviral packaging mix (Cellecta, Inc.), according to the manufacturer’s protocol. High titer viral particles were used to transduce hESCs in serum-free conditions and antibiotic selection of transduced hESCs was performed without MEF feeder cells, using mTeSR1 stem cell media (Stemcell Tech. Inc.). The efficiency of *RB1* or *P53* knockdown and production of cMYC or cMYC (T58A) were determined by Western Blot after antibiotic selection using the following antibodies: anti-RB1 (Cell Signaling, clone 4H1, Cat# 9309), anti-P53 (Santa Cruz, clone DO-1, Cat# Sc-126), anti-MYC (Cell Signaling, clone D84C12, Cat# 5605), anti-HA (Cell Signaling, clone 6E2, Cat#2367), and anti-GAPDH (Cell Signaling, clone 14C10, Cat# 2118). All antibodies for Western Blot were used at a 1:500 working dilution.

### Immunohistochemistry

Living cells in culture were directly fixed in 4% paraformaldehyde for 25 min, followed with 15 min permeabilization in 1% Triton X-100. Histology on tissues from mice was performed on paraffin-embedded or frozen sections from xenografted tumors and corresponding normal tissues, as previously described (10). Tissues were either fixed overnight in 10% buffered formalin and transferred to 70% ethanol, followed by paraffin embedding, or snap frozen in O.C.T medium (Fisher Scientific, Pittsburgh, PA) and fixed in 10% buffered formalin, followed by paraffin embedding. For immunofluorescence, cells or tissue sections were immunostained with antibodies and counterstained with 4,6-diamidino-2-phenylindole (DAPI). Adjacent sections stained with H&E were used for comparison.

Antibodies used for immunostaining or western blot experiments are as follows: anti-CGRP (Sigma, Clone CD8, Cat# c9487), anti-ASCL1 (Sigma, clone 2D9, Cat# SAB1403577), anti-NCAM1/CD56 (R&D systems, cat# AF2408), anti-Ki67 (Cell Signaling, clone D2H10, Cat# 9027, anti-MYC (Abcam, clone Y69, Cat# ab32072), and anti-NEUROD1 (Sigma-Aldrich, clone 3H8, Cat# WH004760M1). All antibodies were used at a 1:150 working dilution.

### Fluorescent activated cell sorting (FACS)

FACS to detect definitive endoderm cells was performed using APC anti-c-KIT (Invitrogen, Cat# CD11705), PE anti-human CXCR4 (Biolegend, Clone 12G5, Cat# 306506) and APC anti-EpCAM (Invitrogen, Clone G8.8, Cat# 17-5791-80) at 1:100 dilution. Cells were incubated with primary antibodies for 30 minutes at 4°C, then washed and suspended in 0.1% BSA/PBS buffer. PE and APC filters were then used to detect cells double positive for KIT and CXCR4, or EpCAM and CXCR4 by signal intensity gating. FACS with anti-CGRP antibody (Abcam, clone 4902, Cat# ab81887) was used to detect CGRP+ cells. Cells were first incubated with anti-human CGRP antibody for 30 minutes at room temperature followed with incubation (1:1000) of secondary antibody conjugated with R-phycoerythrin (PE) for 30 min at room temperature. Then cells were washed and suspended in 0.1% BSA/PBS buffer. PE filter was then used to separate cells into CGRP+ and CGRP-sub-groups by signal intensity gating. Negative controls stained with isotype controls were performed with sample measurements. Cells were sorted on a BD FACSAria II and data were analyzed in FlowJo.

### Xenograft formation

0.3-0.5 × 10^6^ hESC-derived lung cells or PNECs (at day 55 with or without 30 days of prior exposure to 10 µM DAPT alone or to 10 µM DAPT and 1 µM doxycycline to reduce *P53* and *RB1*, and also induce *MYC* expression) were injected subcutaneously or under the renal capsule of 6-8 weeks old, female NOD.Cg-*Prkdc*^*scid*^ *Il2rg*^*tm1WjI*^/SzJ (NSG) mice (Jackson Laboratory, Bar Harbor, Maine) (11). Doxycycline diet began the day after injection (Teklad; 625ppm); tumor development was monitored at least 2-3 times weekly. When animals became moribund or tumor size reached protocol limitations, mice were sacrificed, necropsy performed, and tumors harvested for further histological or molecular study.

### Single cell sequencing and data analysis

Single-cell capture, reverse transcription, cell lysis, and library preparation were carried out by the WCM Epigenomics Core Facility using Chromium Single Cell 3’ v3 kits according to the manufacturer’s protocol (10x Genomics, USA). Single cell suspenions of RUES2-dervied RPM and RPM (T58A) day 50 cultures were prepared by incubating wells in 0.05% Trypsin/0.02% EDTA for 10-15 min. Digestions were then quenched with excess 0.1% BSA/PBS buffer and passed through 40µm mesh strainer. EGFP+ singlets were sorted at the WCM Flow Cytometry Core Facility on a BD Influx or Sony MA900 cell sorter. Each RUES2-dervied RPM and RPM(T58A) day 50 culture pools were isolated from 3 wells of a 12-well plate across two biological replicates, targeting ∼50,000 viable, EGFP+ viable cells per replicate. Cell counts were then adjusted to 6000-9000 cells/50uL in PBS by Trypan Blue exclusion to achieve an estimated capture of 4000-5000 cells. Sequencing was performed by the WCM Genomics Resources Core Facility on a HiSeq 2500 (Illumina, paired-end protocol with 26 base pairs for read 1 and 98 bases for read 2). Single cell RNA sequencing from RP cultures were obtained from previous experiments (5). Transcript abundance was quantified by mapping sequencing reads to a custom human transcriptome reference (GENCODE v42, hg38) that included EGFP and was performed using simpleaf/alevin-fry (12).

Single-cell analyses, including quality filtering, integration, clustering, differential gene expression and reduced-dimensionality visualization, were performed using the Seurat package in R as described in the package SCTransform tutorial (Version 4.3.0) (13). Briefly, cells to be included in the analysis were required to have at least 4000 and at most 45000 unique molecular identifiers (UMIs) and expressed between 1000 and 7,500 unique genes. In addition, cells were excluded if more than 10% of the determined RNA sequences mapped to mitochondrial genes or if they were classified as doublets by scds (14). In total, 7,728 cells from two RP culture samples, 1,875 cells from RPM cultures, 1,562 cells from RPM (T58A) cultures, 3,401 cells from RPM tumors and 3,463 cells from RPM(T58A) tumors passed these filters for quality. Samples were integrated using Harmony with the top 15 principal components and was run with up to 50 iterations of the algorithm (15). Dimensionality reduction was achieved by creating a Uniform Manifold Approximation and Projection (UMAP) of the first 15 dimensions of harmony integration. Clustering resolution was set at 0.1 for the integrated dataset, resulting in three clusters. Subclustering was conducted in a similar manner and involved re-integration of cells from the neuroendocrine cluster and used identical parameters. Cell type annotation was called using the Azimuth functionality within the Seurat package followed by manual inspection, and enrichment analysis was conducted using enrichR (16). A lung atlas containing 584,884 cells was obtained for Azimuth mediated reference mapping from the HubMap portal (14, 17-24).

### Bulk RNA sequencing of the xenograft tumors

Total RNA from the primary and metastatic tumors was isolated using Direct-zol RNA Miniprep kit (Zymo Research). Following RNA isolation, total RNA integrity was checked using a 2100 Bioanalyzer (Agilent Technologies). RNA concentrations were measured using the NanoDrop system (Thermo Fisher Scientific). Preparation of RNA sample library and RNA-seq were performed by the Genomics Core Laboratory at Weill Cornell Medicine or Northwestern University. Messenger RNA was prepared using TruSeq Stranded mRNA Sample Library Preparation kit (Illumina), according to the manufacturer’s instructions. The normalized libraries were pooled and sequenced on Illumina Novaseq 6000 sequencer with pair-end 50 cycles. Sequencing adapters were trimmed, and quality metrics were generated using fastp (25). Transcript abundance was quantified by mapping sequencing reads to a reference transcriptome (GENCODE v42, hg38) using Salmon (26).

### Differential gene expression and enrichment analysis

Identification of differentially expressed genes involved adjusting for sample identity and *cMYC* expression status when appropriate, and was achieved using model-based analysis of single cell transcriptomics (MAST) (27). Gene set enrichment analysis was conducted by mapping differential expression results to hallmark pathways using clusterProfiler (version 4.0) (28, 29). Cell-level enrichment scores for top 20 cluster markers were previously identified (30) using the AddModuleScore functionality of Seurat. Patterns of *MYC* and *NEUROD1* co-expression were limited to cells in which at least one *MYC* and one *NEUROD1* transcript were detected post-processing.

### Deconvolution of bulk RNA sequencing xenografts into SCLC subtypes

Processed single cell RNA-seq data of primary SCLC were obtained from the cellxgene portal (30). Bulk RNA sequencing data for human SCLC tumors (Reads Per Kilobase per Million mapped reads; RPKM) were obtained from cBioPortal. Subtype annotations for 45 patients were obtained from previously published classifications (31). Deconvolution of bulk RNA sequencing data was achieved through BayesPrism (version 2.0) (32). Briefly, single cell transcriptomes from SCLC-A, SCLC-N, SCLC-P and lung adenocarcinoma (LUAD; “NSCLC”) samples were annotated as tumor cells. For deconvolution, aggregated tumor annotations were provided as cell type labels and specific tumor types (e.g. SCLC-A) were provided as tumor *states*. BayesPrism received inputs from single cell annotations and associated count matrix, xenograft and primary tumor bulk RNA-sequencing data and was run with outlier.cut=0.01 and outlier.fraction=0.1. Analysis of resultant cell type fractions was limited to SCLC-A, SCLC-N and SCLC-P. Estimated cell type fractions were converted to proportion metrics within the three SCLC subtypes and visualized for concordance with published annotations.

### Statistical analyses

Group sample sizes for all in vivo experiments were estimated based off of our previous studies, with at least 5 animals per experimental arm being included for purposes of statistical testing (3-5, 33, 34). For mouse experiments, animals were randomly assigned to each group and no animals were excluded from the analyses. All statistical tests are 2-sided. No adjustments were made for multiple comparisons. The relevant investigators (HJC, EEG and KZ) were blinded to experimental allocations among different experimental arms. For all parametric statistical analyses, data were determined to be normally distributed by the D’Agostino-Pearson test. For comparison between experimental and control groups at a specific time point or tissue site in Figures 2, 4, 5, and S1, 2-sided Student t-tests, one-way or two-way ANOVA test, Fisher’s exact tests and two-sided Kolmogorov–Smirnov test were used in GraphPad Prism. Calculated p-values at or below 2.2×10^−16^ were reported as “p < 2.2×10^−16^”, in line with statistical reporting in R.

## Results

### Generation of hESC-derived PNEC-like cells that express transgenes encoding CMYC

We used lentivirus vectors encoding doxycycline (DOX)-inducible mRNAs for wildtype (WT) and mutant (T58A) human cMYC (*see Methods*) to infect previously described hESCs from the RUES2 cell line carrying transgenes encoding DOX-inducible short hairpin RNAs (shRNAs) complementary to *TP53* and *RB1*. For simplicity of presentation, we refer to the latter RUES2 cells as RP cells, and the RP cells containing DOX-inducible WT or T58A mutant cMYC as RPM and RPM(T58A) cells throughout the text and figures of this paper.

After growth of infected cells and differentiation into lung cell progenitors (LPs and LCs; **Figure 1A**), we induced pulmonary neuroendocrine cells (PNECs) by inhibition of NOTCH signaling with DAPT, as previously reported (5). The ability of the RPM and RPM(T58A) cells to produce MYC protein was documented by Western Blotting of cell extracts prepared 72 hours after addition of DOX (**Figure 1B**). As expected, both cell lines also contained dramatically reduced amounts of TP53 and RB1 proteins as a result of DOX-induction of shRNAs specific for these tumor suppressor mRNAs (**Figure 1B**).

**Figure 1.**
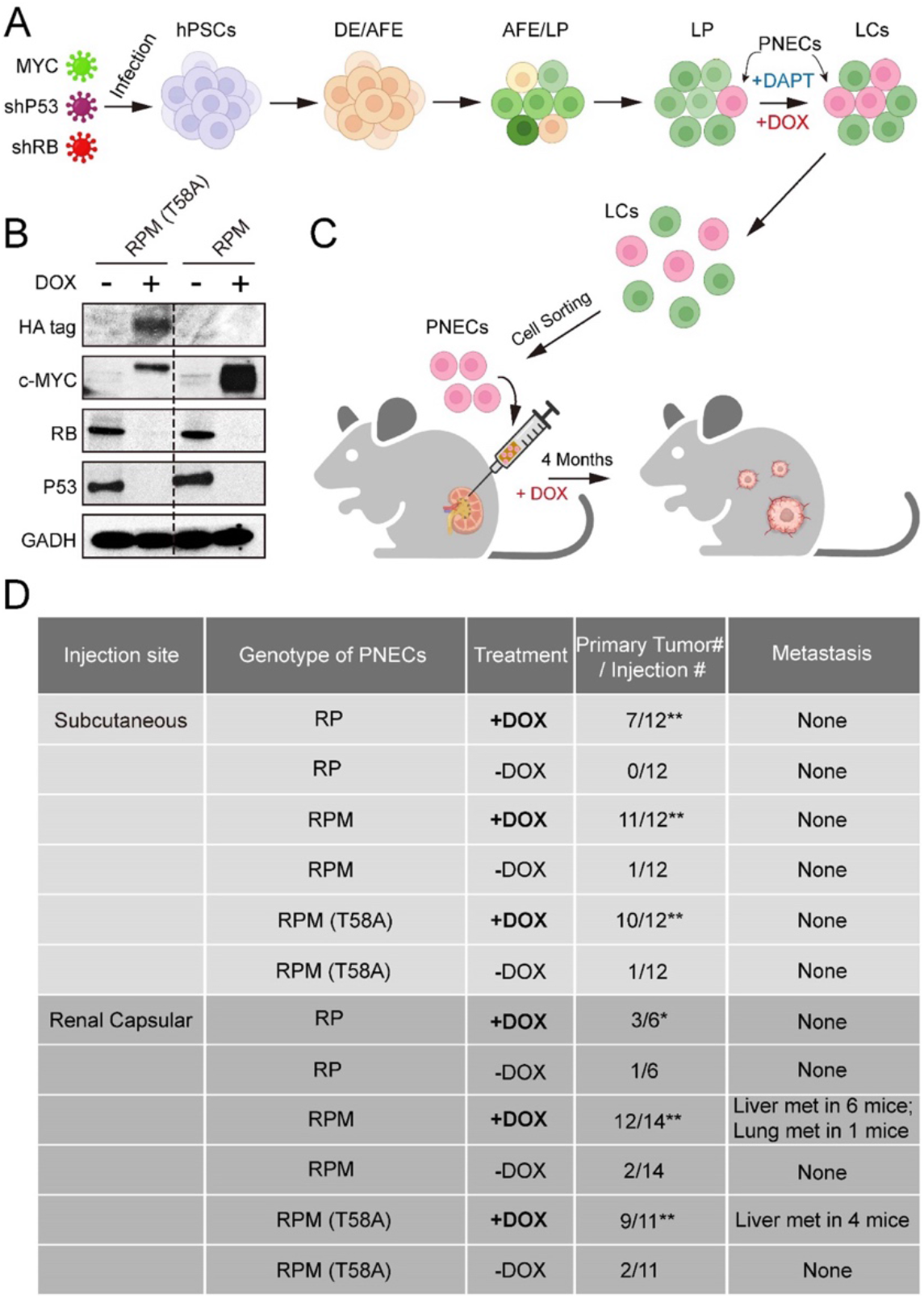
Generation and characterization of hESC-derived lung cells and formation of xenografts with RPM cells. **A)** Schematic of the protocol used to generate PNECs by stepwise differentiation of hPSCs to form: definitive endoderm (DE), day 3; anterior foregut endoderm (AFE), day 6; and lung progenitor cells (LPs) days 15 - 25. LPs were further differentiated into the types of lung cells (LCs) found in mature human lung parenchyma and airway epithelium, days 25 - 55. DAPT (10 μM) encourages the formation of PNECs, and addition of doxycycline (1 μM; DOX) induces expression of shRNAs against *RB1* and *TP53* mRNAs, as well as expression of cMYC or cMYC (T58A), as described in the text. **B)** Western blot of extracts of RUES2 LCs at day 25 of differentiation protocol treated with DOX (1 μM for 72hrs); cells unexposed to DOX served as negative expression controls. Apparent differences in cMYC protein levels may be attributable to the HA-tagged version of cMYC (T58A), which migrates slightly slower that wildtype cMYC protein. **C)** Schematic representation of tumorigenesis experiments comparing injection sites (renal capsule or subcutaneous), DOX treatment (+/-DOX diet), and genotypes (see *Methods* for additional details). Total numbers of animals are 6-7 per experimental arm with 2 injection sites per mouse (right and left flank). Renal capsule injections were performed on a single kidney. Transgenic lines of RUES2 hESCs were differentiated and grown in DAPT (10 μM) from days 25 - 55. At day 55, PNECs were separated from other LCs by sorting for PE+ CGRP-expressing cells (*see Methods*). PNECs were then injected either subcutaneously or into the renal capsular space in NOG mice, half of which then received DOX in their feed as described in Materials and Methods. **D)** Table summary of experiments with xenografted mice, indicating the number of animals that developed visible tumors (≥250 mm^3^ in volume) at the site of injection or the number of visible metastases in the liver or lung. *, P < 0.05; **, P < 0.01 by Fisher’s test to denote significant differences between mice that did and did not receive DOX diet. As before, abbreviations for cell lines are: RP = shRB1 + shTP53; RPM = shRB1 + shTP53 + WT cMYC; RPM (T58A) = shRB1 + shTP53 + cMYC (T58A).

### RUES2-derived PNECs that express WT or T58A cMYC show enhanced oncogenicity as xenografts in the renal capsule of immunodeficient mice

To determine whether production of WT or mutant MYC proteins renders PNECs derived from RPM lines more oncogenic, we sorted CGRP-positive RPM and RPM (T58A) cells from cultures of LCs treated with DAPT and injected the cells subcutaneously or into the renal capsule of immunodeficient mice (the NOG strain: NOD/Shi-scid/IL-2Rγnull) as depicted in **Figure 1A**. The mice were then maintained on diets with or without DOX (*see Methods*). Three to four months later, subcutaneous growths larger than 1000mm^3^ or renal engraftments greater than 100mm^3^ and grossly metastatic lesions in the liver and lungs were tabulated, as summarized in **Figures 1C** and **1D**. As previously reported, PNECs from cultures of RP lines of RUES2 cells produced small benign lesions subcutaneously without invasion or metastasis. Similar findings were observed when these cells were injected into the renal capsule (**Figure 1D**).

### Growth of RUES2-derived RPM tumors are DOX-dependent

Thus far, our data suggest that at least some RUES2-derived, PNEC-like cells in which expression of two tumor suppressor genes, *RB1* and *TP53*, is restricted, RP cells, can be transformed into aggressive, metastatic cells by the addition of transgenes encoding WT or mutant cMYC protein (hereafter referred to as “RPM” or “RPM (T58A)” cells). However, we had not directly compared the relative growth rates of these engineered cell types.

We therefore engrafted immunocompromised mice subcutaneously with equivalent numbers of RP, RPM, or RPM (T58A) cells and monitored tumor growth rates. We noted that the volumes of RPM and RPM (T58A)-derived tumors quickly exceeded the volumes of RP-derived tumors (**Figure 2A**). The rapid growth of RPM tumors was dependent on continuous provision of DOX in the diet. If animals on a DOX-free diet were engrafted subcutaneously with RPM or RPM (T58A) cells, they failed to produce tumors (**Figure 2B**; *left*). However, tumor growth was observed in some mice engrafted with such cells if the animals were placed on a DOX-containing diet up to one month after injection (**Figure 2B**; *right*). These findings suggest that at least some of the RPM cells remained viable and responsive to the oncogenic effects of depleting RB1 and TP53 and producing cMYC.

**Figure 2.**
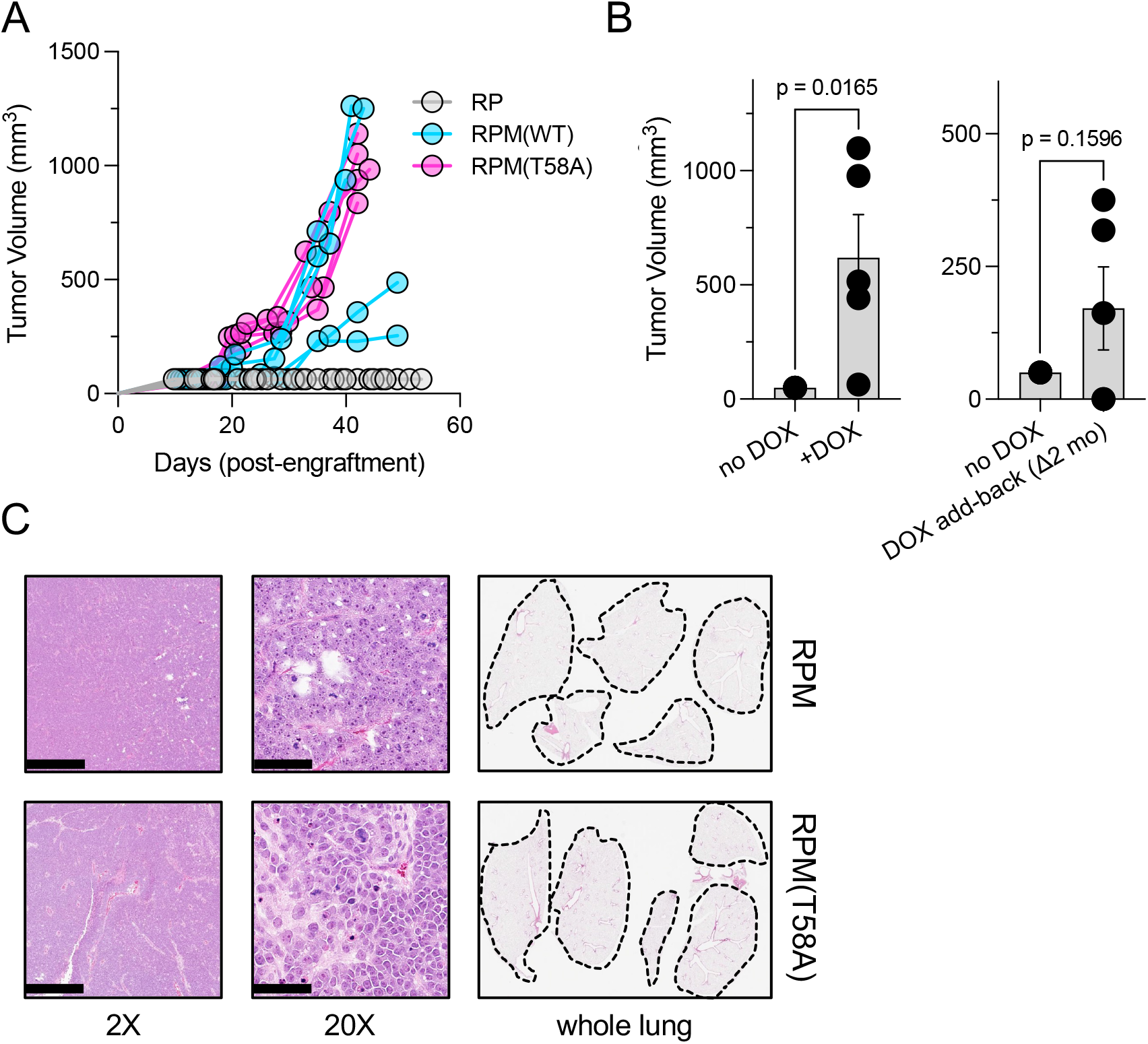
Dox-dependent growth of RUES2-derived RPM and RPM (T58A) subcutaneous tumors. **A)** Subcutaneous engraftment of 10^6^ viable cells at ∼ day 50 of RUES2 lung differentiation protocol from indicated genotypes. Cells were cultured in the presence of media containing 1 μM DOX and 10 μM DAPT from days 25 – 55. All immunocompromised mice were maintained on DOX diet throughout this study; n=5/arm. **B)** *Left*: subcutaneous engraftment of 10^6^ viable RPM tumor cells from the first passage (mouse-to-mouse passage) into immunocompromised mice (NSG) maintained on DOX or normal chow; n=5 animals per arm with single flank engraftments, +/-standard error of the mean (SEM). \*\**P* < 0.05. *Right:* after one month on normal chow, mice were placed on DOX chow (DOX “add-back”); tumor volumes were measured one month later (2 months on study). Paired t test p values are shown for each comparison. **C)** Representative H&E tumor histology from RPM or RPM (T58A) engraftments; 2X (scalebar 1mm) and 20X (scalebar 100um). Representative whole lung sagittal sections showed no evidence of gross metastasis from flank injections at study endpoints; n=3 analyzed per genotype. Dashed lines shown for boundaries of lung lobes.

The rapid growth and features of RPM and RPM (T58A) tumors in these experiments were histologically consistent with SCLC. Critically, subcutaneous injection did not produce metastastic disease to the liver (*not shown*) nor was there evidence of gross metastases to the lungs in any of the mice examined. (**Figure 2C**). These findings suggest that SCLC-like tumors composed of RUES2-derived RPM cells grow rapidly after subcutaneous implantation, but may require a different microenvironment, such as the vascular rich, highly perfused renal capsule (35, 36), providing a more hospitable environment for metastatic seeding.

### Inoculation of the renal capsule facilitates metastasis of the RUES2-derived RPM tumors

RPM tumors, as compared to RP tumors, were notable for increased vascularization, larger volume and invasion into surrounding endothelium (**Figures 3A** to **C**). RPM and RPM (T58A) PNECs injected into the renal capsule of immunocompromised mice also displayed frequent metastasis to the liver (**Figure 3D**). In animals administered DOX, histological examinations showed that approximately half developed metastases in distant organs, including the liver or lung. One of the mice developed metastatic tumors in both liver and lung sites. (**Figure 1D**). No metastases were observed in the bone, brain, or lymph nodes. Metastatic lesions were characterized as having similar proliferative character and expression of cMYC, as well as comparable expression of the neuronal markers NCAM (CD56) and NEUROD1 compared to their paired primary counterparts (**Figures 3E** and **3F**). In contrast, ASCL1 expression was absent in RPM and RPM (T58A) tumors at both primary and metastatic sites (**Figure 3F**). Meanwhile, the majority of cells in RP tumors showed positive ASCL1 expression but lacked NEUROD1 expression (**Figure 3G**). These findings suggest that MYC expression may facilitate the transition of SCLC tumors from the classical ASCL1-driven subtype to the NEUROD1 variant. Thus, both the site of implantation of cells and the addition of a driver oncogene were required to produce vigorous growth and metastatic spread of SCLC-like tumors, and the metastatic lesions retained histological and immunohistochemical features of the primary tumors.

**Figure 3.**
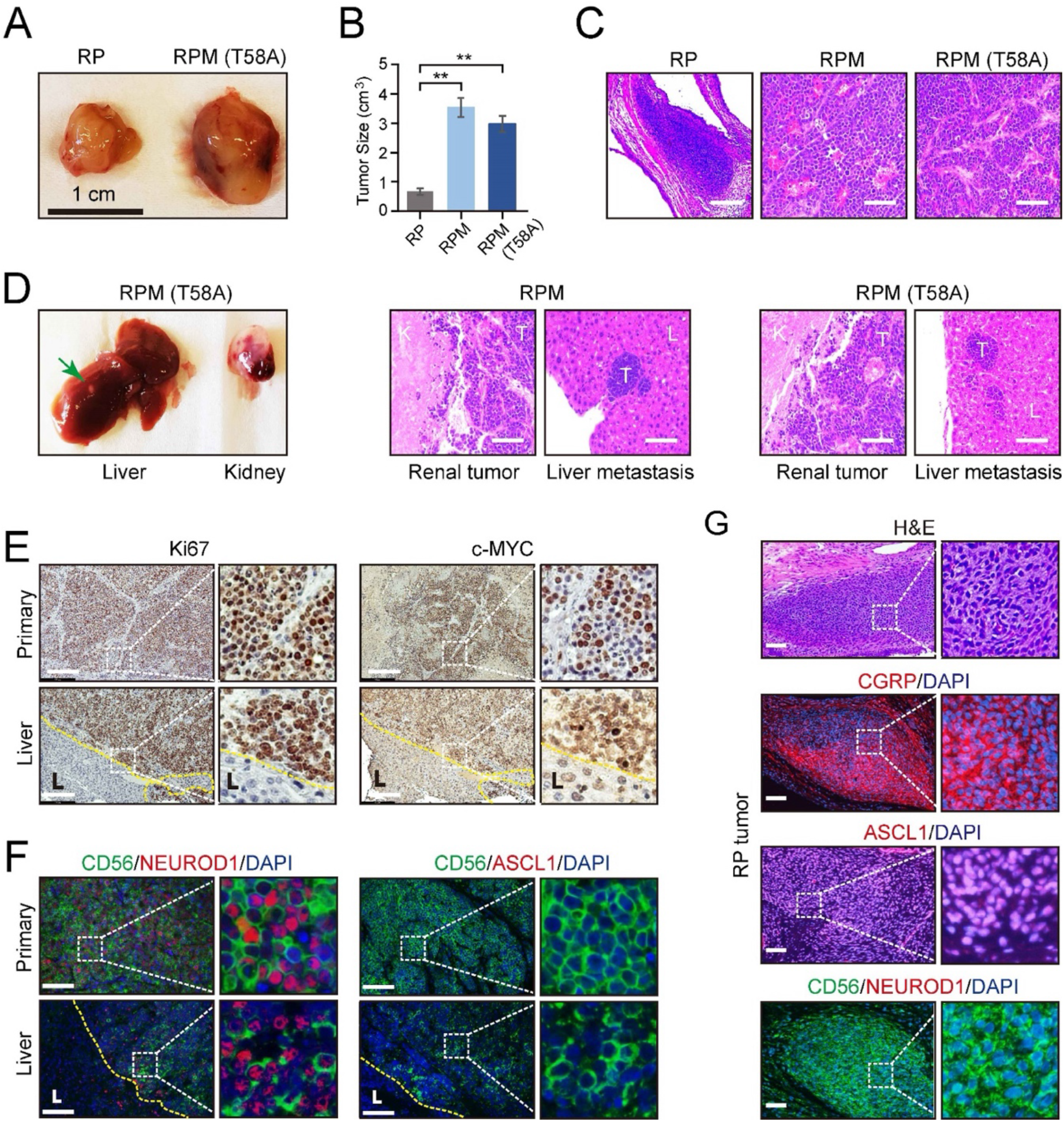
Effects of cMYC on the gross and histopathological appearance of xenografts grown in immunodeficient mice fed with DOX for 3-4 months. **A)** Subcutaneous xenografts formed with hESC-derived PNECs with RP, RPM or RPM (T58A) genotypes. Photographs of representative tumors formed with cells of the indicated genotypes are shown for RP and RPM; indicated scale of 1cm. **B)** Quantification of the tumor sizes and paired comparisons for a secondary *in vivo* experiment; n=5 animals per arm with single subcutaneous engraftmetns; **P<0.01. **C)** H&E staining of the indicated tumors from panel *B*. **D)** Gross and histologic pathology of renal capsular xenografts and liver metastases formed with the RPM or RPM (T58A) cells. Left panels, gross appearance of representative tumors within the liver (metastasis) or kidney (primary); right panels, H&E staining of primary and metastatic tumors (T) formed in kidney (K) and liver (L) in the RPM (left) or RPM (T58A) model. **E)** Immunostaining of tumor samples from *D*. Samples of primary peri-renal tumors and hepatic metastases in mice injected with hESC-derived PNECs programmed to reduce levels of *TP53* and *RB1* mRNA and to express wild type MYC were stained with antisera for Ki67 (*left*) or MYC (*right*). **F)** Immunofluorescence staining for neuroendocrine markers ASCL1, NEUROD1 and CD56 from sections in *E* of the RPM tumors. **G)** Immunofluorescence staining for neuroendocrine markers, CGRP, ASCL1, NEUROD1 and CD56 in the RP tumors. H&E staining serves as the bright-field comparison of the indicated tumors.

### RUES2-derived RPM tumors most closely resemble the NEUROD1-high (SCLC-N) subtype of SCLC

To better understand the characteristics of RUES2-derived RPM tumors, we performed single-cell RNA-sequencing on culture-derived (“2D”) and tumor-derived RPM and RPM (T58A) viable EGFP+ cells. By integrating previously published RUES2-derived RP single cell data (5), we observed three dominant cell-type groups in transcriptional space, using reference-mapping, enrichment analysis and manual curation to identify cell lineage likelihood: a basal-like cluster marked by expression of *KRT19*, a fibroblast-like cluster marked by expression of *COL1A1*, and the most prominent cluster of cells appearing to have features of neuroendocrine differentiation, marked by expression of the neuroendocrine lineage defining transcription factor *ASCL1* (**Figure 4A** and **Supplemental Table 1**). Comparing the recently described SCLC subtype markers (31) across these datasets showed that cells within the neuroendocrine differentiation cluster cells expressed *ASCL1* and *NEUROD1*, whereas *POU2F3* RNA was not detected. Levels of *YAP1* RNA were variable and were not co-expressed with either *ASCL1* or *NEUROD1* (**Figure 4B**). *YAP1* expression was mostly attributed to fibroblast- and basal-like cells that are often observed in long-term cultures of differentiated hESCs (**Figure 4B**) (4, 37-39).

**Figure 4.**
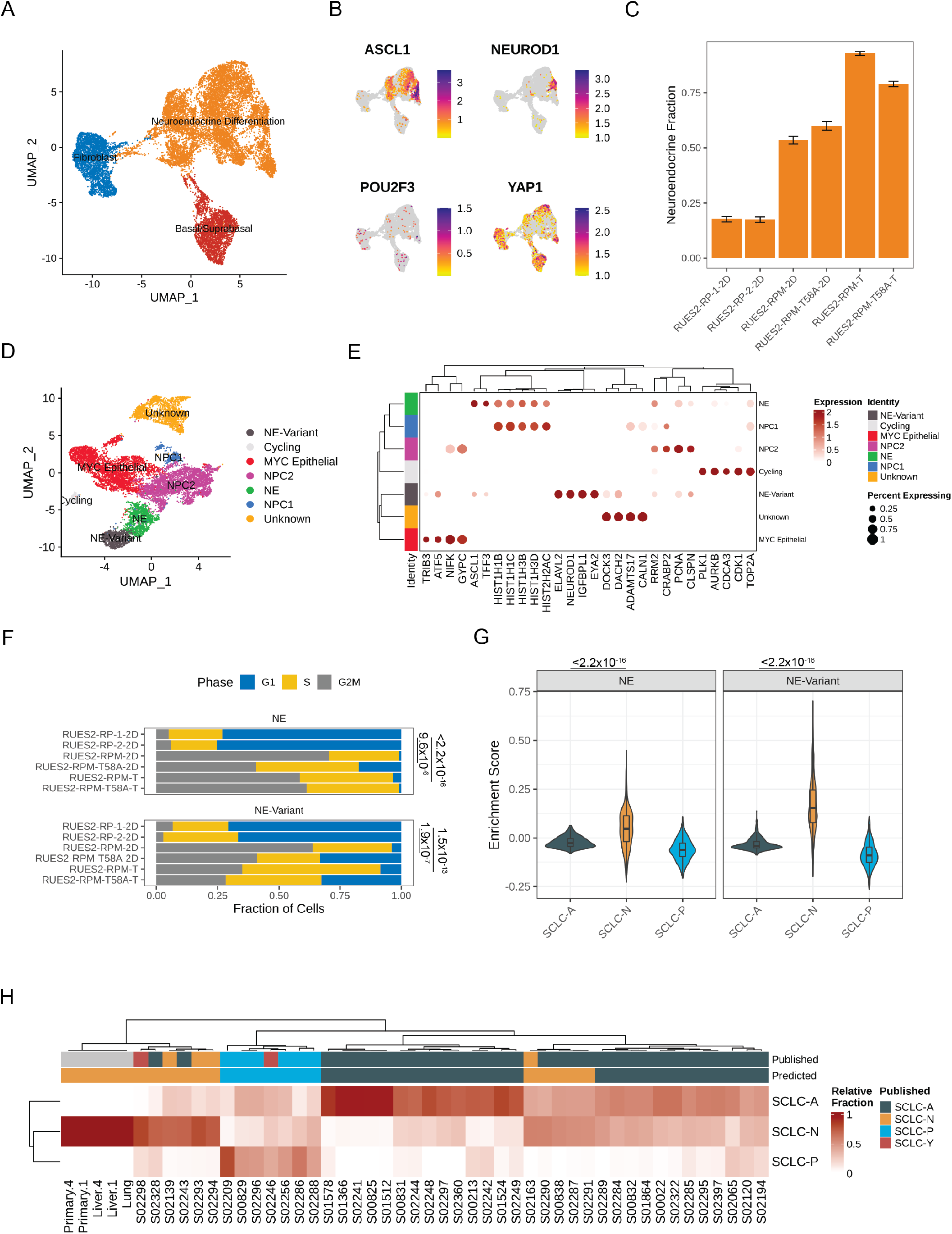
Single cell and bulk RNA profiling of RUES2-derived RPM tumors and comparison with primary human SCLC. **A)** UMAP projection of cultured RP samples from Chen et al (2019) and RPM samples [*this study*] with 3 major cellular lineages annotated by color. **B)** Expression of SCLC subtype markers across dataset in *A*. **C)** Neuroendocrine fraction of cells by sample ID. **D)** Subclustering of the “neuroendocrine differentiation” cluster from *A*. **E)** Dot plot of differential cluster markers in the subclustering analysis from *E*. **F)** Cell cycle evaluation of NE and NE-Variant clusters indicated by color and fraction of cells. **G)** SCLC subtype enrichment scores from Chan et al (2021); cluster markers in NE and NE-Variant cells from RPM tumors. **H)** Bulk RNA-sequencing subtype estimation based on Chan et al (2021) SCLC subtypes from RPM primary or metstatic tumors as compared to published primary SCLC data. Published labels were obtained from Rudin et al (2019). Bulk patient RNA-sequencing data (reads per kilobase per million mapped reads (RPKM)) were compared to bulk RNA-sequencing of select RPM tumors and metastatic samples. Primary tumor and liver metastases were obtained from two pairs (animals #1 and #4) of mice engrafted with RUES2-derived RPM tumor cells into the renal capsule and grossly macro-dissected during necropsy.

Next, we asked whether the over-expression of *MYC* in RPM cells was associated with increased neuroendocrine differentiation when compared to cell lines that did not contain c*MYC* transgenes (i.e. RP cells). One notable difference between our prior work and this study was our attempt to increase the tumor cell component of the RPM and RPM (T58A) samples by sorting cells for high EGFP expression. EGFP is expressed in *cis* with the *RB1* targeting shRNA; however, EGFP is not expressed uniformly across all cells following the differentiation protocol, suggesting some cells may silence the integrated vector. We performed differential cluster abundance analysis after accounting for the fraction of cells that were EGFP*+* during cell sorting. These results indicated that RPM cell lines (WT or T58A) were associated with a two-fold increase in the neuroendocrine compartment when compared with RP cells (p < 2.2×10^-16^; **Figure 4C** and **Supplemental Figures 1A** and **1B**). Similarly, RPM tumors had a three-fold enrichment of cells expressing neuroendocrine markers compared to RP cell lines (p < 2.2×10^-16^). Notably, we found that RPM tumors and cells expressed *MYC*, not *MYCN* or *MYCL* at high levels (**Supplemental Figure 1C**). Moreover, *EGFP* expression was strongly correlated with *MYC* (Pearson r = 0.45, p < 2.2×10-16) and not *MYCN* (Pearson r = 0.00043, p = 0.97) nor *MYCL* (Pearson r = −0.031, p = 0.002). Together, these results suggest that the over-expression of either WT or T58A MYC yielded a larger neuroendocrine compartment.

By sub-clustering the neuroendocrine differentiation group, we could clearly identify *ASCL1*-high and *NEUROD1*-high compartments, as well as several other groups of cycling cells that did not cleanly fit into any lung lineage cell identity (**Figure 4D, Supplemental Table 2**). We conducted differential gene expression to further characterize the *NEUROD1*-high (abbreviated “NE-variant”) and *ASCL1*-high (abbreivated “NE”) clusters observed with RPM cells (from cell lines and tumors) and found that the NE-variant cluster was associated with increased expression of genes highly expressed in variant small cell lung cancer (*NEUROD1, SCG3, IGFBPL1, SSTR2* and c*MYC)* (**Supplemental Figure 1D, Supplemental Table 3**). In contrast, the NE (or “NE-classic”) cluster was linked to heightened expression of a variety of genes ranging from histones (*HISTH1B, HISTH1D*) to redox processes (*MT1E, MT2A*) (**Figure 4E**). Gene set enrichment analysis of these results in the NE-variant cluster revealed increased expression of genes associated with activation of epithelial-mesenchymal transition (EMT; p = 6.81×10^-3^) and pancreatic beta cell signaling (p = 1.01×10^-3^). However, this cluster was also linked to lower levels of oxidative phosphorylation (p = 1.07×10^-25^), possibly suggesting these sub-groups of cells are biologically different (**Supplemental Figure 1E**). Further examination of the *ASCL1*-high and *NEUROD1*-high clusters revealed that, across both clusters, RPM and RPM (T58A) cells were associated with higher expression of genes associated with cells in dividing phases of the cell cycle than were observed RP cultures. We compared the proportion of G2M cells between culture-derived RPM and RP models. This revealed that RPM cells were associated with a 10- and 11-fold increase in the NE and NE-variant clusters respectively (NE p = 9.6×10^-6^, NE-Variant p = 1.9×10^-7^) with a relative reduction in cells in the G1 phase (**Figure 4F**). Similar trends were observed from data in human tumors (NE p < 2.2×10^-16^, NE-Variant p = 1.5×10^-13^, **Figure 4F**).

Given the striking increase in proliferation in cMYC producing cells, we sought to understand the association between c*MYC* and *ASCL1* or *NEUROD1* expression. We computed the cellular density of co-expression patterns of cells that express c*MYC* in addition to either *ASCL1* or *NEUROD1*. This identified that the joint density was highest for cells that co-express c*MYC* and *NEUROD1* and was localized to the *NEUROD1*-high cluster (**Supplemental Figure 1F**). In comparison, c*MYC* and *ASCL1* co-expression patterns were heterogenous, and found in the *ASCL1*-high cluster as well as in cells that could not be annotated with confidence. Similar to previously reported by other groups, we found that *ASCL1* and *MYC* expression are negatively correlated (Pearson r = −0.11, p = 1.5×10^-4^). Moreover, modest correlation between c*MYC* and *NEUROD1* expression was identified in our single cell data (Pearson r = 0.17, p = 3.7×10^-4^, **Supplemental Figure 1G**). Given that single cell sequencing experiments suffer from a high dropout rate, we attempted to validate c*MYC* and *NEUROD1* co-expression patterns in bulk primary and metastatic RPM tumors. Although we couldn’t determine co-expression within the same cell using bulk gene expression data, we identified an extremely strong correlation between the expression of these two genes (Pearson r = 0.93, p = 0.02, **Supplemental Figure 1H**). Taken together, we show that the over-expression of *MYC* (WT or T58A) may be linked to an SCLC-N phenotype.

Upon identification of the *ASCL1*- and *NEUROD1*-high clusters, we sought to further characterize how closely they resemble SCLC tumors transcriptionally using a two-pronged approach. First, we overlayed SCLC-A, SCLC-N and SCLC-P signatures (30) from our single cell data and calculated subtype enrichment scores. Enrichment patterns in RPM and RPM (T58A) xenografts revealed that both *ASCL1*- and *NEUROD1*-high clusters expressed SCLC-N-associated genes more frequently than SCLC-A and SCLC-P-associated counterparts (p < 2.2×10^-16^, **Figure 4G**). In contrast, RP cell cultures more closely resembled cells from tumors with an SCLC-A phenotype (p < 2.2×10^-16^, **Supplemental Figure 1I**). Second, we identified the transcriptional identity of RPM xenografts and paired metastases by cellular deconvolution. Bulk gene expression of RPM tumors along with human SCLC biopsies (40) were deconvolved into SCLC subtypes. This identified relative cellular proportions of SCLC subtypes in bulk RNA-seq data. Overall, we found high concordance with published subtype annotations and those predicted by our analysis (**Figure 4H**) (31). Moreover, we found that all RPM xenografts strongly exhibited an SCLC-N transcriptional program. Taken together, these data suggest that over-expression of wild type or mutant *MYC* alters the transcriptional identity of RP cells, encouraging neuroendocrine differentiation, and may assist with the transformation from a SCLC-A to SCLC-N phenotype.

## Discussion

In this report, we show that the addition of an efficiently expressed transgene encoding normal or mutant human cMYC can convert weakly tumorigenic human PNEC cells, derived from a human ESC line and depleted of tumor suppressors RB1 and TP53, into highly malignant, metastatic SCLC-like cancers after implantation into the renal capsule of immunodeficient mice.

Experimental models for understanding the origins, behaviors, and control of human cancers have been developed in numerous ways for more than a century. Some of the models depend on producing tumors in animals or transforming the behavior of cells growing in culture with viruses, chemical agents, genetic changes, or other perturbations, or by deriving cells from human cancers and growing them in animals or in tissue culture. The choice of these approaches have been influenced by the goals of the research, the type of cancer under study, and available technologies.

Because SCLC is a highly aggressive cancer with a relatively stereotypic genotype that is generally recalcitrant to therapeutic strategies that have not changed appreciably in the past thirty years, we have been attempting to generate a model for SCLC in which the cancer cells are human in origin, derived from the most commonly affected pulmonary cell lineage (neuroendocrine), and equipped with the major and most common genotypic and phenotypic changes characteristic of SCLC.

The most obvious use of these cells (the RPM lines described here) is for the evaluation of the efficacy of novel therapeutic agents acting directly on tumor cells during growth in culture or during growth as xenografts in immunosuppressed mice. However, our attempts to conduct therapeutic studies with the RPM lines have not been sufficiently extensive or rigorous to warrant presentation here. In addition, treatment strategies that involve host elements other than the tumor itself, such as immune cells or other features of the tumor micro-environment, might require an experimental platform more complex that those that we have constructed.

Our choice of cMYC to serve as an oncogenic driver in hESC-derived PNECs was based on both the frequent finding of high levels of *cMYC* RNA in human SCLC and the use of Myc over-expression strategies to generate aggressive mouse models of SCLC (7, 8, 41-43). However, not all human SCLC tumors have abundant *cMYC* RNA, and other genes, including other members of the MYC gene family, have been implicated in the pathogenesis of SCLC. So it will be interesting to compare the ability of transgenes encoding other MYC proteins, such as *NMYC* and *LMYC*, as well as other oncoproteins (both those implicated and not implicated in human SCLC carcinogenesis) to convert RP cells into malignant tumors and to produce metastatic phenotypes, as we have observed here with RPM cells. In particular, it may be important to determine whether hESC-derived SCLC cells with varied genotypes are liable to metastasize to specific sites such as the brain and bone, which are commonly affected in patients with SCLC, but were not sites for metatastic growth in the experiments reported here. Finally, the role of other tumor suppressor genes, such as *PTEN* – implicated in the development of mouse models of histological transformation of LUAD to SCLC that we have recently reported (44) – should also be examined in this human ESC-derived model.

We have used transcriptional profiling as a means to compare human ESC-derived SCLC grown from RPM cells with SCLC tumor samples from patients. Other aspects of tumor cell behavior – including pathophysiological features, responses to therapeutic strategies, and the appearance of molecular, metabolic, and immunological markers – might also provide means to determine the degree of similarity between this novel stem cell-derived model and genuine human neoplasms.

Transcriptional profiling has also allowed us to conclude that the hESC-derived tumors arising from RPM RUES2 cells belong to the SCLC-N subtype, one of the major subtypes of SCLC now recognized by several laboratories and clinicians studying SCLC. It is unclear whether SCLC-N truly represents a subtype of SCLC with distinct outcomes and therapeutic sensitivities (45), or rather a snapshop from a continuum in tumor evolution driven [principally] by *MYC* towards an aggressive, recalcitrant disease (46, 47). Future studies of stem cell-derived SCLC should consider ways to generate tumors that model the other common subtypes including SCLC-P and SCLC-I (“inflamed” SCLC) (45). This might be achieved by altering the cancer genotype, by using a differentiation scheme that produces physiologically varied precursor cells, or by employing a wider variety of hESCs to develop SCLCs.

Finally, we note that in the experiments reported here metastatic spread was observed from RPM tumors growing initially in the renal capsule but not from primary subcutaneous tumors initiated with the same cell culture. It may prove important to test other sites of primary tumor growth with RPM cells from the RUES2 line or with other SCLC models, to determine the degree to which metastatic spread depends on the site(s) of growth of the primary tumor and to identify specific factors that promote such spread.

## Author Contributions

H.J.C., E.E.G., K.Z. and A.T. performed wet lab experiments. Y.S. and O.E. performed computational analyses, including single cell RNA-sequencing data analysis. All authors participated in the design and interpretation of some or all experiments. H.V., H.J.C., and E.E.G. wrote the manuscript, and all authors suggested editorial changes. H.V. and H.J.C. conceived the study and recruited the collaborating partners.

## Acknowledgements

We thank Oksana Mashadova and Sukanya Goswami in the Varmus Laboratory for technical support, Arun Unni and John Ferrarone in the Varmus Laboratory and Jonathan Chen in the Chen Laboratory for useful advice. Supported by awards to H.V. and O.E. from the US Department of Defense (LC160136) and the U.S. National Cancer Institute (U01CA224326), funds from the Meyer Cancer Center, Weill Cornell Medicine (to H.V.), and a K99/R00 NIH Pathway to Independence Award (4R00CA226353), US Department of Defense (RA220012) and a Lung Cancer Research Foundation (LCRF) Pilot Project Award (to H.J.C.).

## SUPPLEMENTAL FIGURES

**Supplemental Figure 1.**
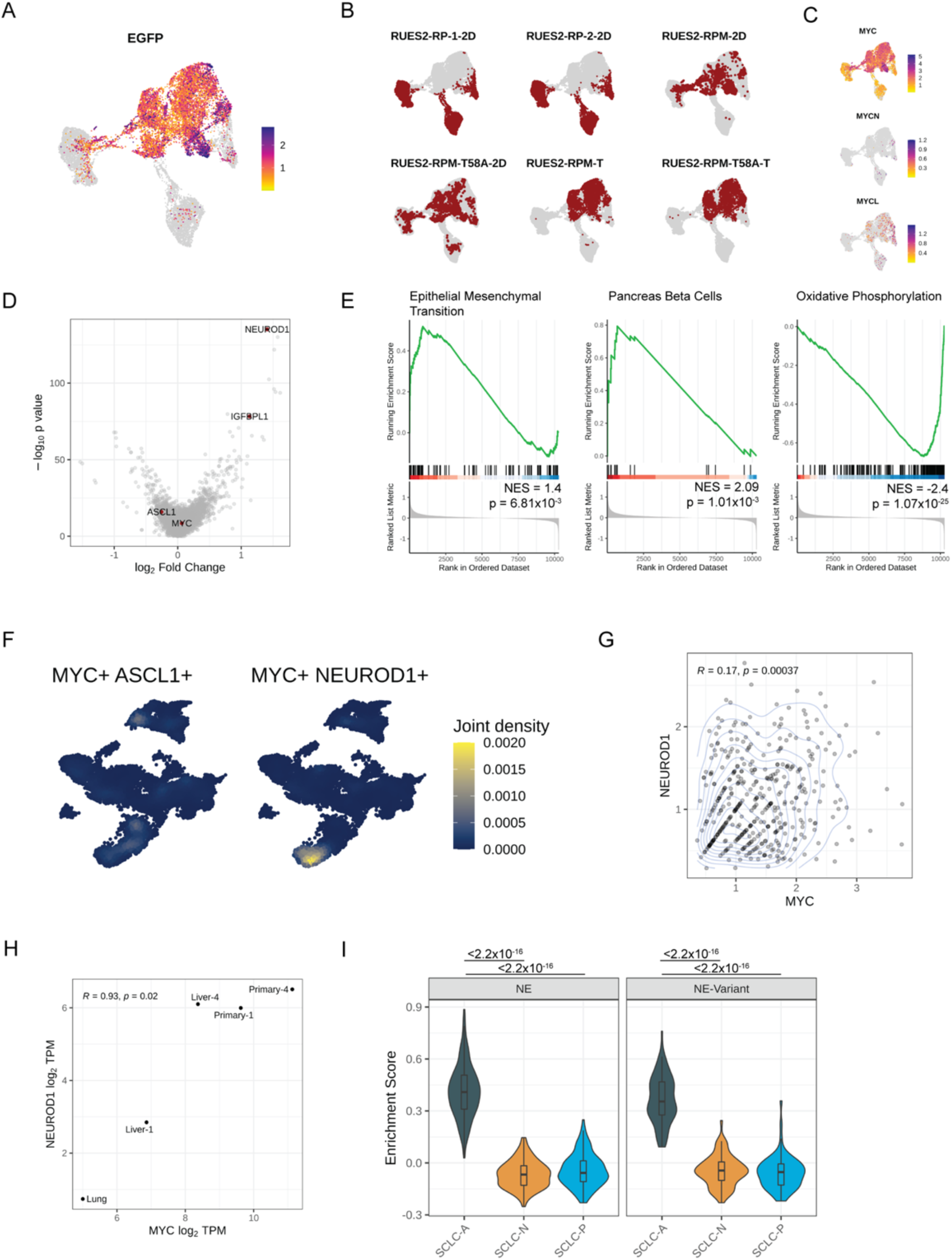
Gene expression and characterization of RUES2-derived tumors. **A)** *EGFP* expression across the full dataset integrating RP and RPM cells in addition to RPM tumor cells. **B**) Individual sample IDs and relative contribution (weighted; red) to initial clustering distribution for the 6 sample sets used in this combined analysis. **C)** *MYC, MYCN* and *MYCL* expression across the full dataset **D)** Differential gene expression (DEG) between NE-Variant and NE cluster. Select genes are called out in red. **E)** Select gene set enrichment analysis for top 3 differentially expressed programs between *NEUROD1*-high (NE-variant) and *ASCL1*-high (NE) subsets of cells from only RPM tumor cells; shown as normalized enrichment scores (NES). Positive enrichment is greater activation in the NE-Variant subset (i.e. the *NEUROD1*+ subset), whereas negative enrichment is greater in the NE subset. **F)** Cellular density of co-expression patterns for *ASCL1* and *NEUROD1* across the neuroendocrine differentiation subset. **G)** Scatter density plot of *MYC* and *NEUROD1* expression across all cells in the neuroendocrine differentiation cluster. **H)** Plot of *MYC* and *NEUROD1* expression in bulk expression data derived from RPM tumors; data are shown as log2 of transcripts per million (TPM). **I)** Enrichment score for SCLC subtype markers in NE and NE-Variant cells from RP cell cultures (2D), demonstrating preferably enrichment of the SCLC-A subtype in the absence of over-expression of a *MYC* transgene.

**Supplemental Figure 2.**
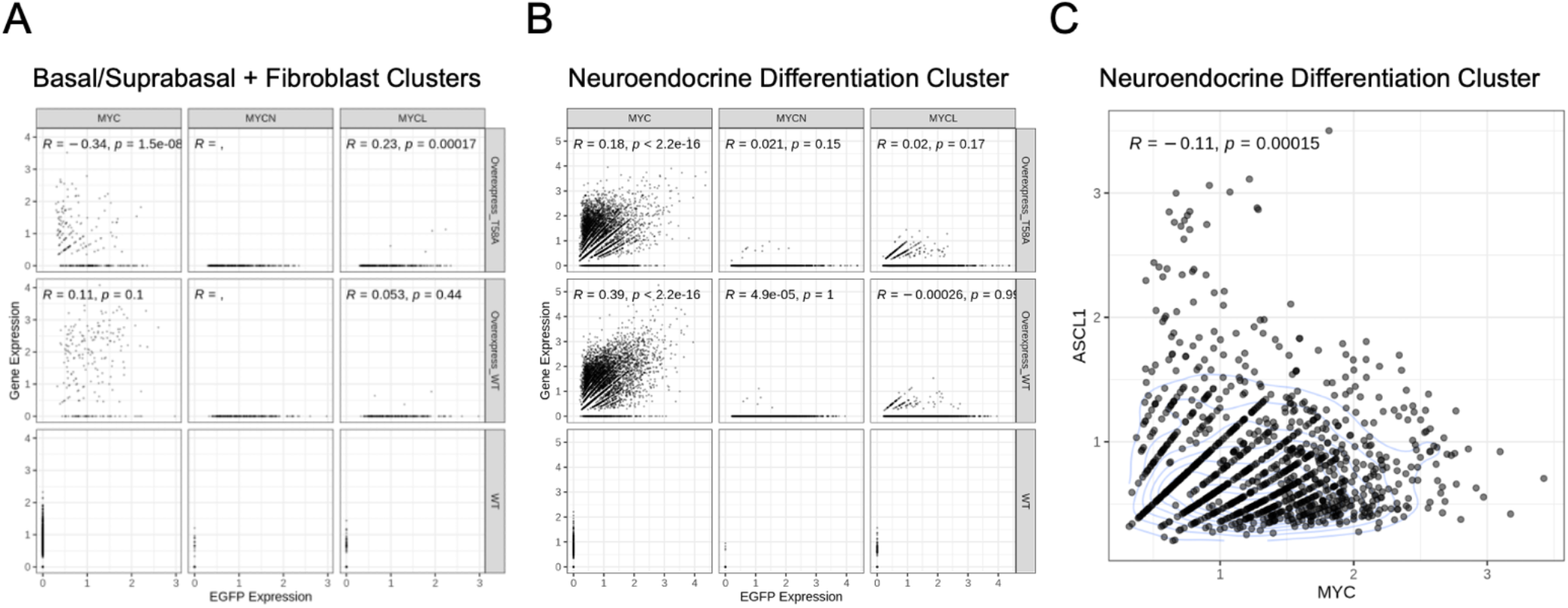
MYC family member expression correlates in RUES2-derived RPM tumors. **A**) Expression correlation of *EGFP* for *MYC, MYCN* or *MYCL1* from the neuroendocrine differentiation cluster identified in Figure 4A. **B)** Similar to A, but now for the neuroendocrine differentiation cluster. Scatter plots depict correlation in NE and non-NE clusters. **C)** Expression correlation of *ASCL1* and *MYC* in the neuroendocrine differentiation cluster. All sample pools were included in this correlation. Each data point is a single cell. Correlation (*R* value) and significance (*P* value) shown for each individual comparison.

